# *In vivo* brain activity imaging of interactively locomoting mice

**DOI:** 10.1101/203422

**Authors:** Shigenori Inagaki, Masakazu Agetsuma, Shinya Ohara, Toshio Iijima, Tetsuichi Wazawa, Yoshiyuki Arai, Takeharu Nagai

## Abstract

Electrophysiological field potential dynamics have been widely used to investigate brain functions and related psychiatric disorders. Conversely, however, various technical limitations of conventional recording methods have limited its applicability to freely moving subjects, especially when they are in a group and socially interacting with each other. Here, we propose a new method to overcome these technical limitations by introducing a bioluminescent voltage indicator called LOTUS-V. Using our simple and fiber-free recording method, named “SNIPA,” we succeeded in capturing brain activity in freely-locomotive mice, without the need for complicated instruments. This novel method further allowed us to simultaneously record from multiple independently-locomotive animals that were interacting with one another. Further, we successfully demonstrated that the primary visual cortex was activated during the interaction. This methodology will further facilitate a wide range of studies in neurobiology and psychiatry.

## Introduction

Brain functions and related psychiatric disorders have been investigated by recording electrophysiological field potentials^1^. Electroencephalography (EEG) measures whole brain wide activity, while local field potential recording detects the dynamics of more localized micro-networks of the neurons^2^. More recently, the development of fluorescence-based organic dyes and genetically encoded indicators to detect neuronal activity has proven useful for brain-wide and local micro-network level recordings^1,3–9^. Both electrophysiological and fluorescence-based techniques have been further developed to record freely moving animals, via electrical or optic fibers connected to them. This allows the detection of brain activity during more complicated behavioral tasks^1,10–13^.

However, there are still several technical limitations associated with the use of these conventional methods. One limitation is the requirement of complicated instruments and special knowledge of electrophysiology or optics. Another limitation is that the subject must be head-fixed or connected to the cable (electrical or optical), which limits the application of these techniques to freely moving subjects, especially when recording from a group of multiple animals interacting with each other. Therefore, investigating neural networks regulating social behaviors and related psychiatric diseases, required a fiber-free detection system, which was suitable for multiple animals. Moreover, recording techniques that are simply and easy to use are ideal for further application to “brain-activity based” high-throughput screening systems, such as the search for drug effects or genetic factors.

Here we report a novel fiber-free recording method using a recently described bioluminescence-based voltage indicator “LOTUS-V”^14^. Our method enables simultaneous recording from multiple animals interacting with each other, using a very simple optical setup. In comparison to recently reported methods for *in vivo* voltage imaging from freely moving animals, our method was advantageous for studying neural networks regulating social behaviors, and has the potential to be applied for further high-throughput screening systems that are based on *in vivo* activity recording.

## Results

### Voltage imaging in neurons using LOTUS-V

LOTUS-V consists of a voltage-sensing domain (VSD) from voltage-sensing phosphatase (Ci-VSP)^15^, NLuc (i.e., a cyan-emitting luciferase that is approximately 150 times brighter than Renilla or firefly luciferases)^16^, and Venus (i.e., a yellow fluorescent protein)^17^. An increase of the membrane voltage causes a structural change of the VSD, enhancing Förster resonance energy transfer (FRET) between the NLuc and Venus, which consequently, decreases the NLuc signal and increases the Venus signal (**Supplementary Fig. 1**). We previously demonstrated that LOTUS-V is useful to monitor voltage changes in HEK293T cultured single cells and moving cardiomyocyte aggregates. Furthermore, we verified the advantages of LOTUS-V in long-term imaging and the robustness of the signal reliability in a moving specimen^14^.

To validate the efficacy of LOTUS-V in voltage imaging of neurons, we first expressed it in primary cultures of rat hippocampal neurons, and investigated signal changes during patch clamp recording (**Fig. 1** **and Supplementary Fig. 2**). An intense bioluminescence was observed from a single neuron on addition of furimazine, i.e., a substrate for the NLuc moiety in LOTUS-V (**Fig. 1a** **and Supplementary Fig. 1b**). Thus we could record the signal using a 1-kHz frame rate (**Fig. 1c-d** **and Supplementary Fig. 2**). As previously observed in HEK293T cells^14^, LOTUS-V had wide-range detectability in neurons (**Fig. 1b**; −120 mV to +80 mV), suggesting that it could detect both subthreshold activity and an action potential. The time constants of the voltage response (**Fig. 1c**) were like those of a conventional indicator, ArclightQ239 (ref. 18, 19). During neuronal firing, a significant signal increase of LOTUS-V was observed (**Fig. 1d**; 0.80±0.29% increase [mean ± SE] compared to baseline; n=6 cells; p <0.05, one sample t-test). In additiion, the calculation of the emission intensity ratio from two channels rather than a single channel was advantageous to obtaining a larger signal change (**Supplementary Fig. 2a**). Even a single trial or averaging a small number of trials could produce a clear signal change (**Supplementary Fig. 2b**). The majority of recorded cells independently showed significant signal changes (**Supplementary Fig. 2c**). These results substantially confirmed the efficacy of LOTUS-V in voltage imaging of neurons, which further suggested that it could reliably report *in vivo* brain activity (i.e., electrophysiological field potential dynamics derived from neuronal population). A previous study also demonstrated that LOTUS-V was advantageous to analyze voltage changes in a mass of cardiomyocyte cells^14^.

**Figure 1.**
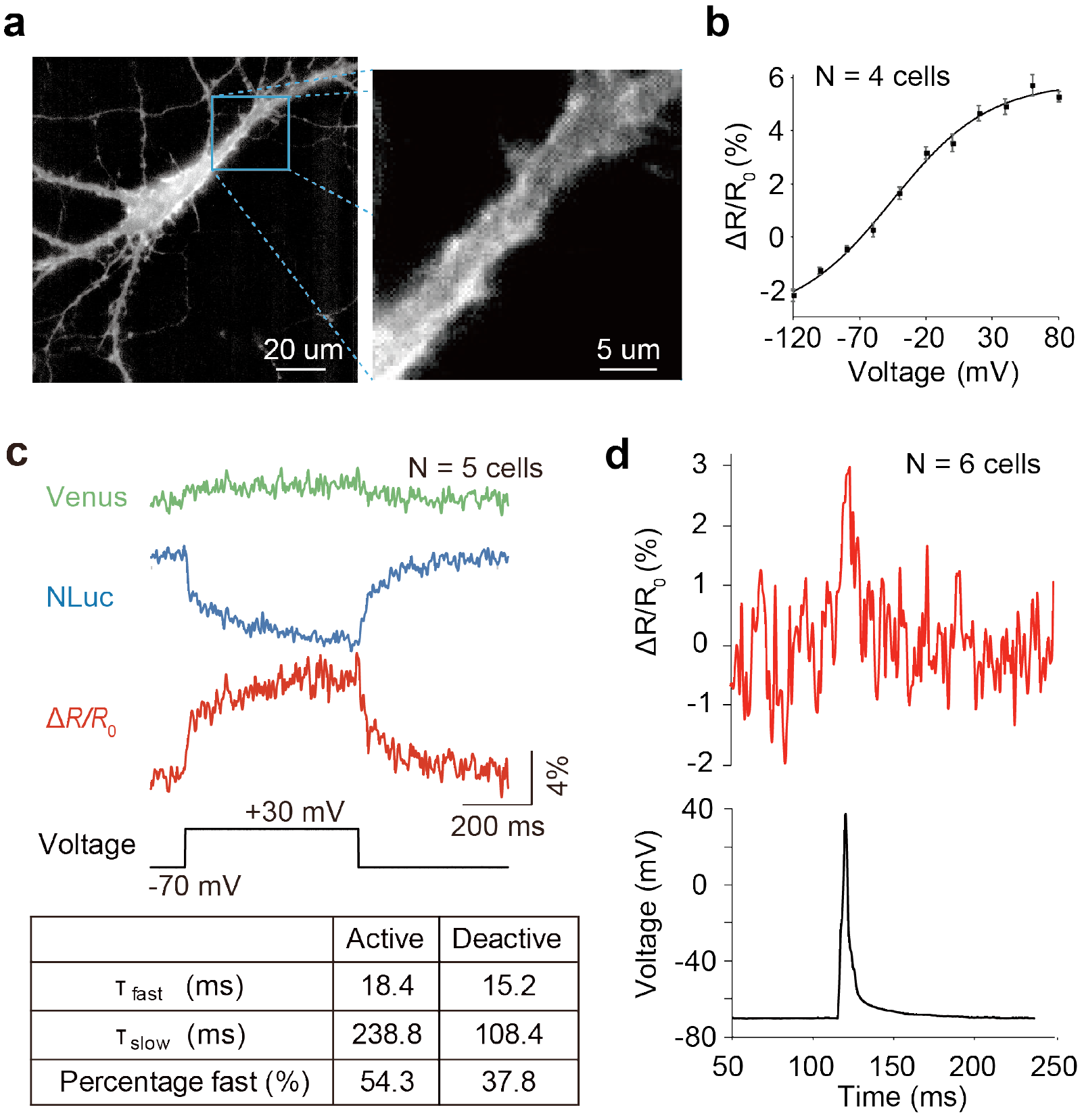
Electrophysiological characterization of LOTUS-V in hippocampal neurons. (**a**; left) A representative bioluminescence image of a cultured hippocampal neuron expressing LOTUS-V. (**a**; right) An expanded image of the region within the square on the image on the left. (**b**) Plot of fractional Δ*R*/*R*_0_ versus voltage changes (n=4 cells). The Δ*R*/*R*_0_ from −120 mV to +80 mV was 5.3±0.3%. The effective valence (Z) was 0.7, while the V_1/2_ was −45.5 mV. The plot was fitted using a Boltzmann function. (**c**) The Venus and NLuc signals (Δ*L*/*L*_0_), and their ratio (Δ*R*/*R*_0_), in response to voltage changes from the holding voltage (−70 mV to +30 mV; n=5 cells). (**c**; table) The fast and slow components, and their fraction of time constant. The activation and deactivation curves of AR/R_0_ were fitted using a two-component exponential equation. (**d**; upper) Action potential waveform of Δ*R*/*R*_0_ and (**d**; lower) electrophysiology (n=6 cells). The imaging frame rate was 1 kHz. Error bars indicate mean ± standard error.

### LOTUS-V can report brain activity in an awake and head-fixed mouse

We first tested the efficacy of LOTUS-V for *in vivo* imaging of brain activity from an awake mouse, using a head-fixed system. LOTUS-V was locally expressed in the primary visual cortex (V1) using the adeno-associated virus (AAV) gene expression system, and labelled a population of local neurons in the V1 (**Fig. 2a-c**). Since it was already well confirmed that neural activity in the V1 was correlated with the locomotion of mice, even in the absence of visual input or in the dark^20–22^, we tested whether LOTUS-V signal could reflect such a locomotion-dependent increase of neural activity.

**Figure 2.**
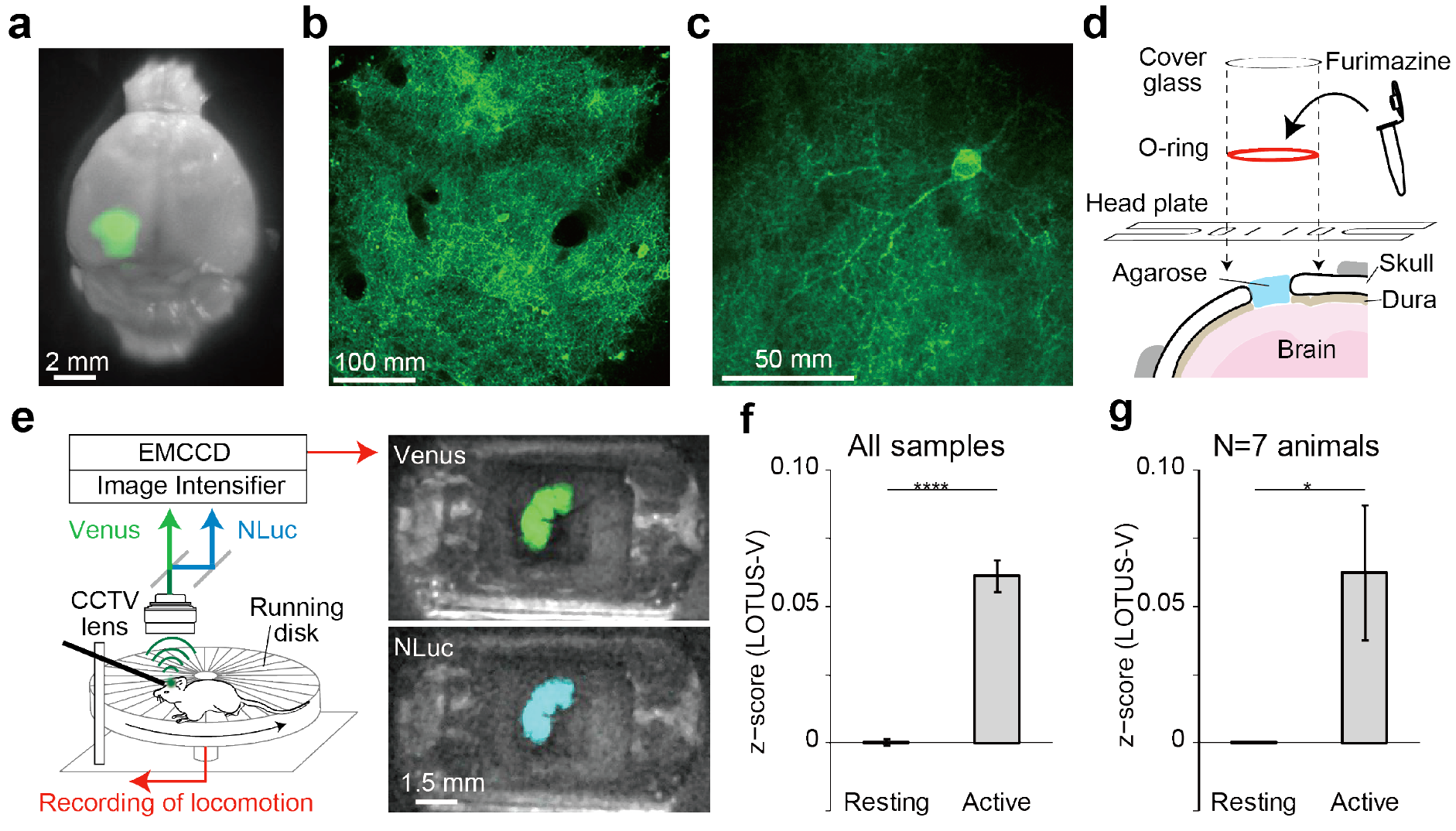
Imaging of a head-fixed mouse with LOTUS-V. (**a**) Venus fluorescence in the primary visual cortex (V1) of a paraformaldehyde-fixed brain. (**b-c**) Examples of two-photon fluorescence images of V1 area containing LOTUS-V-expressing (**b**) neurons and (**c**) a single V1 neuron. (**d**) Schematic drawing of the prepared cranial window. The headplate and o-ring were glued over the cranial window. The furimazine solution was enclosed in the o-ring bath by a cover glass. (**e**; left) Schematic drawing of the imaging setup for a head-fixed mouse. A mouse was placed on a running disk while the head was fixed to a bar via the headplate. The locomotion velocity was recorded through a rotary encoder during imaging. LOTUS-V bioluminescence from the V1 was collected through a CCTV lens. NLuc and Venus emissions were separated by image splitting optics. These emissions were acquired using an image intensifier unit, which can amplify the signal up to 10^6^ times, and recorded with an EMCCD camera in the dark condition. (**e**; right) Overlaid images of bright field and bioluminescence in NLuc and Venus channels acquired by this system. (**f-g**) Plots of z-normalized Δ*R*/*R*_0_ in resting (<5 cm/s) and active (>5 cm/s) states of head-fixed mice, using (**f**) all time-points (p <0.0001, Wilcoxon rank sum test; n= 707871; 31653 time-points), or (**g**) average values from each mouse (p <0.05, Wilcoxon signed-rank test; N=7 animals). Error bars indicate mean ± standard error; *, p <0.05; ****, p <0.0001

The ratio change of the LOTUS-V signal in the V1 of head-fixed mice was recorded. Furthermore, the speed of their spontaneous locomotion was simultaneously recorded using the rotation of a running disk placed under the animal (**Fig 2d-e**). Using this system, we confirmed that LOTUS-V signal was significantly changed while mice were moving on the running disk (**Fig. 2f-g**).

Since V1 neurons strongly respond to visual stimulation^3,6,8^, we also investigated the signal change of the LOTUS-V during visual stimulation, to further elucidate its applicability for variable sensory modalities (**Supplementary Fig. 3**). Light illumination for the visual stimulation can be reflected and contaminated in the camera, consequently masking the images. Therefore, we conducted the “dead-time imaging”^14,23^, where the exposure to the camera and visual stimulation during the processes for image readout and accumulated charge clearing on the camera (dead-time) were alternately (not simultaneously) performed (**Supplementary Fig. 3a-b**). LOTUS-V successfully reported an increase in signal, depending on the light intensities (**Supplementary Fig. 3c**), which confirmed its applicability to monitor changes in local brain activity, irrespective of the input modality.

### LOTUS-V can report brain activity from a freely moving mouse

Next, we examined whether LOTUS-V can report brain activity in a freely moving mouse. Conventionally, brain activity imaging from a freely moving animal requires a complicated and invasive imaging system, including multiple optical fibers to excite and detect fluorescent voltage indicators and an additional infrared video camera to detect locomotion^1,11^. In contrast, our method only required an animal to be placed in the cage, with a completely detached CCD camera system fixed above the mouse cage (**Fig. 3a-b**). During the experiments, a strong bioluminescence from the targeted brain region was continuously observed for 1.15−6.77 h (3.10±0.45 h [mean ± SE]; N=16 mice), enabling us to perform long-term imaging. Considering TEMPO^1^, i.e., a method recently developed for fiber-coupling based voltage imaging, can only be performed for approximately 1 h, our method is more advantageous for long-term imaging. Moreover, since our method does not require illumination light, limitations due to laser fluctuation, auto-fluorescence, and bleaching do not have to be considered. Our fiber-free method enables recording from freely moving animals with a very simple optical setup, and was thus, named “SNIPA” (i.e., <u>S</u>imple and <u>N</u>o-fiber-attachment voltage <u>I</u>maging <u>P</u>latform for freely moving <u>A</u>nimals).

**Figure 3.**
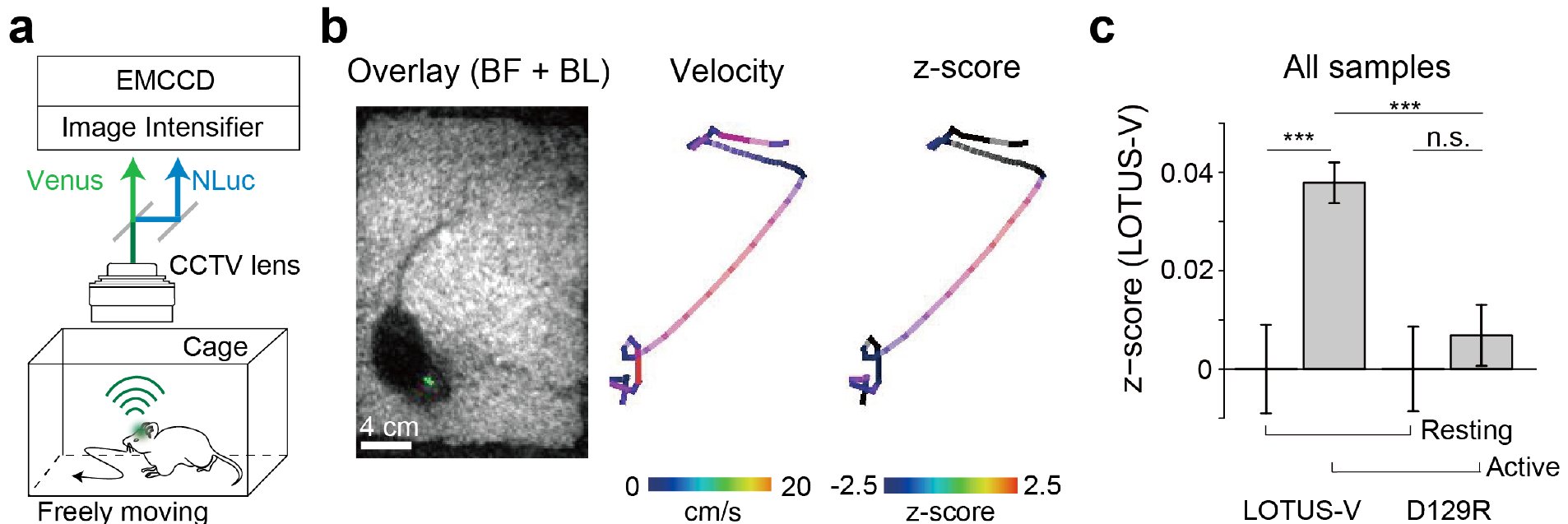
Imaging of a freely moving mouse with LOTUS-V. (**a**) Schematic diagram of imaging of a freely moving mouse. The mouse was placed in its home cage and the LOTUS-V bioluminescence was recorded. (**b**; left) Overlaid image of bright field and LOTUS-V bioluminescence (green). (**b**; middle and right) Pseudo-colored trajectories of mouse locomotion, indicating locomotion velocity (middle) and z-normalized Δ*R*/*R*_0_ (right) (see also **Supplementary Videos 1 and 2**). (**c**) Bar plot of z-normalized Δ*R*/*R*_0_ in the resting (<1 cm/s) and active (>1 cm/s) states of freely moving mice (p <0.0001 for one-way ANOVA with all four categories; resting and active states of LOTUS-V, n= 12277 and 57079 time-points from N=5 animals; resting and active states of LOTUS-V(D129R), n= 13467 and 26573 from N=3; p-values shown in the panel were calculated using a post-hoc Tukey-Kramer test). Similar results were also obtained using a Wilcoxon rank sum test (resting vs active in LOTUS-V, p <0.0001; resting vs active in LOTUS-V(D129R), p >0.05; LOTUS-V (active states) vs LOTUS-V(D129R) (active states), p <0.0001). Time bin, 0.1 s; Error bars indicate mean ± standard; n.s., not significant; ***, p <0.001

The position change of the bioluminescence spot in a freely moving mouse was detected using a particle track analysis program (**Fig. 3b** **and Supplementary Video 1**). We could automatically detect the LOTUS-V signal in the V1 and locomotion velocity (**Fig. 3b** **and Supplementary Video 2**). The z-score (normalized values calculated from Δ*R*/*R*_0_) increased significantly during the active state of a freely moving mouse (**Fig. 3c**; p <0.0001, Wilcoxon rank sum test), in corroboration with previous studies^20–22^. To evaluate whether this signal change reflected a real voltage change or was just an artifact, we compared the results of LOTUS-V expressing animals (N=5 animals) with those expressing voltage-insensitive LOTUS-V(D129R) (N=3 animals). Locomotion did not significantly change the LOTUS-V(D129R) signals. Further, the LOTUS-V signal during the active state was statistically higher than the LOTUS-V(D129R) signal during the active state (**Fig. 3c**; p <0.0001, one-way ANOVA; p <0.001 for the resting vs active states of LOTUS-V; n.s. for the resting vs active states of LOTUS-V(D129R); p <0.001 for LOTUS-V during the active state vs LOTUS-V(D129R) during the active state, post-hoc Tukey-Kramer test). These results certified that SNIPA based on LOTUS-V could be used to analyze neural activity in a freely-moving mouse. A previous study^24^ reported that locomotion-driven change in the neural activity of V1 is mild in the dark or when visual stimulation is absent. It was, however, significantly detected using our system, suggesting the substantial sensitivity of SNIPA for imaging a freely-moving mouse.

### SNIPA can detect brain activity from interactively locomoting mice

Although a technique to measure brain activity during social interaction is important and desirable, the simultaneous targeting of multiple animals that are interactively moving is still very challenging. This is mainly because most current methods rely on fiber-coupled recording for both electrophysiology and fluorescence imaging^1,10–13^. Although a fiber-free method based on bioluminescence was recently reported^25^, this method required very complicated image processing and intensity calibration, and has not been applied to simultaneous imaging of multiple animals. In contrast to these previous methods, here we demonstrated a fiber-free method, which enabled simultaneously recording of multiple mice that were freely interacting with each other (**Fig. 4a** **and Supplementary Video 3**). To capture the mouse body clearly, the inside of the cage was illuminated with a light emitting diode, and bright-field and bioluminescence images were taken altenately^26,27^. Thereafter, we manually tracked the target area of each animal, and measured the locomotion and LOTUS-V signal. Apart from the mouse that was quiet throughout the imaging, and thus, excluded from analysis (see **Online Methods**), the locomotion-driven signal change in the V1 of freely interacting mice was successfully detected (**Fig. 4a** and b). To our knowledge, this is the first report of simultaneous recording of brain activity from three animals that are interacting with one another. Further, this experiment demonstrated that SNIPA can not only be performed in the dark, but also under the illumination.

**Figure 4.**
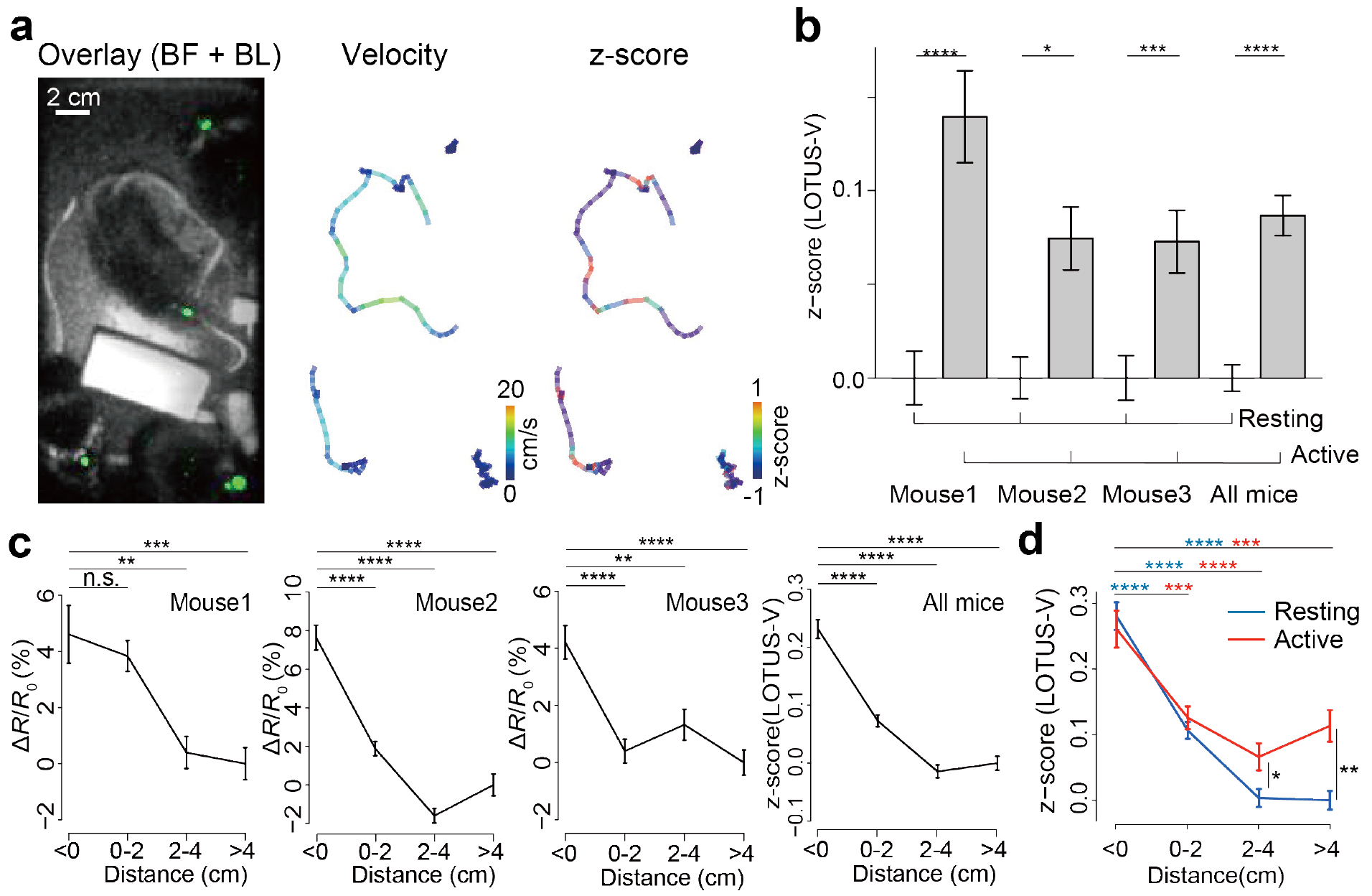
Imaging of interactively locomoting mice. (**a**; left) Overlaid image of bright field and LOTUS-V bioluminescence from multiple mice (green). (**a**; middle and right) Pseudo-colored locomotion trajectory, indicating velocity (middle) and z-normalized Δ*R*/*R*_0_ from each mouse (right) (see also **Supplementary Videos 3 and 4**). (**b**) Bar plots of z-normalized Δ*R*/*R*_0_ during resting (<1 cm/s) and active (>1 cm/s) states of freely-moving multiple mice, demonstrating locomotion-dependent signal increases (Mouse 1, n= 4862 and 1935 time-points; Mouse 2, n= 8051 and 4099; Mouse 3, n= 7062 and 3741; All mice, n= 19975 and 9775). P-values were obtained using a Wilcoxon rank sum test. (**c**) Distance-dependent change in the activity of the primary visual cortex (V1) of interactively locomoting mice. Plots represent Δ*R*/*R*_0_ or z-normalized Δ*R*/*R*_0_ of each distance category (distance between the target mouse nose and other mice, see also **Supplementary Fig. 4**; distance from Mouse 1, n= 665, 2215, 1905, and 2012 time-points for <0, 0-2, 2-4 and >4 cm, respectively; Mouse 2, n= 1892, 4745, 3747, and 1776; Mouse 3, n= 2163, 3462, 2048, and 3139). The distant values (>4 cm) were used as a baseline (*R*_0_) to calculate the Δ*R*/*R*_0_ (for Mouse 1-3) and for the z-normalization (for “all mice”). P-values were obtained using a multiple comparison (Tukey-Kramer) test. (**d**) Comparison of distance-dependent change in V1 activity and the effect of locomotion (resting vs active states). Data from all mice were used (resting state (<1 cm/s, blue), n= 3075, 6769, 5214, and 4914 time-points for <0, 0-2, 2-4, and >4 cm, respectively; active state (>1 cm/s, red), n= 1645, 3653, 2486, and 2008). The Δ*R*/*R*_0_ in the “distant and resting” state was used as the baseline for z-normalization. P-values obtained using a one-way ANOVA with all four categories; p <0.0001 in both states. P-values obtained using multiple comparisons (Tukey-Kramer test) are shown as blue (resting states) or red (active) symbols, while those obtained using the Wilcoxon rank sum test (to compare resting vs active) are shown in black. The time bin was 0.1 s. Error bars indicate mean ± standard error; n.s., not significant; *, p <0.05; **, p <0.01; ***, p <0.001; ****, p <0.0001

More importantly, by systematically analyzing the distance between the mice (**Supplementary Fig. 4**), we demonstrated that V1 activity was significantly higher when each mouse approached the others (**Fig. 4c**). In addition, by separately analyzing the locomotion-dependent and distance-dependent effects (**Fig. 4d** and **Supplementary Fig. 5**), we demonstrated that locomotion did not significantly increase brain activity in the V1 when the distance was small (<2 cm). This suggested that the interaction with other mice may have a stronger or more competitive impact on the V1 compared to simple exploratory locomotion. These results indicated that SNIPA can be a powerful method to investigate novel brain function during social interaction.

## Discussion

Taken together, the results demonstrated that SNIPA is a powerful method to monitor neural activity from a target brain area of freely moving mice. To our knowledge, this is the first report of functional brain activity imaging using a bioluminescent probe in a freely moving animal, without conventional invasive and complicated procedures, such as the insertion of an optical fiber or prism, which can potentially disturb normal brain activity or circuitry. Further, this fiber-free imaging method enabled us to record from interactively locomoting mice, and demonstrated a novel type of V1 activation during the interaction, confirming the importance and usefulness of SNIPA. Since this method uses a genetically encoded indicator, we can perform tissue or cell-type-specific expression of LOTUS-V, as well as whole-brain macroscopic imaging, all of which are widely required in neuroscience research^28^. Furthermore, since optogenetic manipulation is compatible with LOTUS-V imaging^14^, a wireless optogenetics device, which could be combined to SNIPA, may allow us to wirelessly detect and manipulate neural activity in socially interacting animals^29,30^. Overall, SNIPA opens a door to an easy and minimally invasive method to image brain activity simultaneously in multiple socially interactive animals, thus, contributing to a wide range of neuroscience research.

## Acknowledgements

T.N. was supported by a Grant-in-Aid for Scientific Research on Innovative Areas ‘Spying minority in biological phenomena’ (No. 3306) of MEXT (No. 23115003, No. 23115001), Grant-in-Aid for Scientific Research (A) of MEXT (No. 26251018), the JST-SENTAN program, JSPS Core-to-Core Program, A. Advanced Research Networks, the Uehara Memorial Foundation, and the Naito Foundation. S.I. was supported by the “Program for Leading Graduate Schools” of MEXT and a Grant-in-Aid for JSPS Research Fellow (16J00111). M.A was supported by MEXT (No. 15K18341). We thank Hiroshi Hama (RIKEN, BSI) for teaching us neuron culture, Akihiro Yamanaka (Nagoya University, RIEM) for providing us with an AAV purification method, Mitsuhiro Iwaki (RIKEN QBiC) for lending us a C8600-05 GaAsP image intensifier unit, and Robert E. Campbell (University of Alberta, Department of Chemistry) for providing us with the pAAV2-hSyn. We also thank the Bionanophotonics Consortium (BNPC) for assistance in experiments using microscopy.

## Author contributions

T.N. conceived and coordinated the project. S.I., M.A., and T.N. designed the experiments; S.I., S.O., and T.I. performed the electrophysiological experiments.; S.I. constructed *in vivo* imaging system with the support of M.A., T.W., and Y.A.; M.A. performed *in vivo* imaging with the support of S.I.; all the authors analyzed data; S.I., M.A., and T.N. wrote the paper, with contributions from all authors.

## Online Methods

### AAV preparation

For the AAV expression system, the cDNA of LOTUS-V (GenBank accession number; LC061443) and LOTUS-V(D129R) were amplified from pcDNA3-CMV-LOTUS-V and pcDNA3-CMV-LOTUS-V(D129R) respectively, by polymerase chain reaction, using a sense primer, containing a Kozak sequence following a *Bgl*II site, and a reverse primer, containing a *Hin*dIII site and a stop codon. They were then ligated with pAAV2-hSyn^31^ digested by *Bam*HI and *Hin*dIII. DNA sequencing was used to verify all constructs.

The AAV-DJ vector was prepared as described previously with some modifications^32^. Briefly, HEK293T cells (RIKEN BRC Cell Bank, RCB2202) were grown in Dulbecco’s modified Eagle’s medium (Sigma) containing heat-inactivated 10% FBS (Biowest) at 37 °C in 5% CO_2_. Equal amounts of pAAV2, pAAV-DJ^33^, and pHelper (Cell Biolabs, INC.) plasmids were co-transfected using FuGENE_HD_ transfection reagent (Promega), following the manufacturer’s protocol. The cells were trypsinized and centrifuged for 5 min at 1000 rpm at 4 °C, 3 days after infection. The pellet was resuspended in 200 μl HEPES-buffered saline (10 mM HEPES, pH 7.3, containing 150 mM NaCl, 2.5 mM KCl, 1 M MgCl_2_, and 1 M CaCl_2_) and subjected to three freeze-and-thaw cycles. Then, 1 μl of benzonase nuclease was added to each tube, warmed in a water bath at 45 °C for 15 min, and centrifuged for 10 min at 16,000 *g* at 4 °C. The supernatant was transferred into a new Eppendorf tube and centrifuged for 10 min at 16,000 *g* at 4 °C. Thereafter, the supernatant was transferred to a new Eppendorf tube and aliquots were stored at −80 °C until use.

### Animals

All experimental procedures were conducted according to the Institutional Guidance on Animal Experimentation and with permission from the Animal Experiment Committee of Osaka University (authorization number: 3348). C57BL/6JJmsSlc male mice were purchased from Japan SLC, Inc. The mice were housed in the Osaka University Animal Facility and supplied with food and water *ad libitum*.

### Rat hippocampal neuron culture for imaging

Primary cultures of hippocampal neurons and astrocytes were prepared from embryonic day 17 Sprague-Dawley rats. Cells were dissociated in plating medium consisting of Hanks’ Balanced Salt solution (HBSS; Wako), which was supplemented with 1 mM HEPES and 100 U/ml penicillin/streptomycin. The cells were then plated on a poly-L-lysine (Sigma) coated 35-mm dish with a coverslip bottom at a density of 3.5 × 10^4^ cells/12-mm-diameter coverslip. The medium was changed to culture medium constituting of Neuro Basal (Thermo Fisher), supplemented with 2% B27 (Invitrogen) and L-glutamine, 5 h after plating, and the cultures were grown in 5% CO_2_ at 37 °C. On day 7 *in vitro* (DIV-7), half of the medium was replaced with fresh culture medium. On DIV-14, the stock solution containing AAV vector was mixed with culture medium up to 1.0 × 10^10^ TU/ml. After 5 h of incubation, the medium was replaced to fresh culture medium. Cultures were incubated again at 37 °C in 5% CO_2_, and were used for experiments 7-10 days after infection.

### Electrophysiology and photometry

LOTUS-V-expressing hippocampal cultured neurons were subjected to simultaneous patch-clamp and FRET recordings at 7-10 days post-infection. The medium was replaced with a bath solution consisting of HBSS (Gibco) supplemented with 20 mM HEPES (pH 7.2) and 5.5 mM D-Glucose, and the dish was mounted on an Axiovert 200M inverted microscope (Carl Zeiss). Patch-clamp recordings in the whole-cell mode were made using an Axoclamp 200B patch-clamp amplifier, with a capacitive head stage (Axon Instruments), using glass recording electrodes (3-5 MΩ) filled with intracellular solution (i.e., 140 mM potassium gluconate, 5 mM KCl, 1 mM MgCl_2_, 0.1 mM EGTA, 2 mM Mg-ATP, 5 mM HEPES, adjusted to pH 7.2 with KOH). Whole-cell recordings were low-pass-filtered at 1 kHz and digitized at 10 kHz. Data were digitized with a Digi data 1342 digitizer (Axon Instruments) and fed into a computer for offline analysis using AxoClamp 9.0 software (Axon Instruments). Bioluminescence of LOTUS-V was observed by adding 50 μM furimazine (Promega) and recordings were performed at 23 °C. Images of the Venus and NLuc channels were acquired simultaneously using W-VIEW GEMINI image splitting optics (Hamamatsu Photonics), C8600-05 GaAsP image intensifier unit (Hamamatsu Photonics), and MiCAM Ultima-L CMOS camera (Brain Vision). The image splitting optics contained a FF509-FDi01-25×36 dichroic mirror (Semrock) and no emission filters. The optical signal was analyzed offline using BrainVision analyzer software.

### Preparation for *in vivo* voltage imaging

Viruses were injected to C57BL/6JJmsSlc mice at postnatal day P35-40 for *in vivo* experiments. Procedures were conducted as previously described^9^, with some modification. During surgery, mice were anesthetized with isoflurane (initially 2% partial pressure in air, and reduced to 1%). A small circle or a square (~1 mm in diameter) was thinned over the left V1 using a dental drill to mark the site for craniotomy and target the putative monocular region. AAV-DJ vector was injected into the left V1 (2.1 mm lateral to the midline, 0.3 mm rostral to lambda at a depth of 300 μm), over a 15-min period, using a UMP3 microsyringe pump (World Precision Instruments). The total volume of the AAV crude solution was 375 nl. The beveled side of the needle was faced to the left so that viruses could be injected into and cover the V1 area of the left hemisphere.

Mice were anesthetized by isoflurane (1-1.5%), at 3 weeks to 5 months after the virus injection, before the *in vivo* recording. *In vivo* two-photon imaging was performed as previously described^9^ to confirm the expression level of LOTUS-V. Briefly, a titanium headplate was attached to the skull using dental cement, and the cranial window was made over the V1, around the virus injection site (1.5 mm in diameter) for subsequent imaging. The brain surface was covered with 4% agarose gel dissolved in HEPES-buffered saline (10 mM HEPES, pH 7.3, containing 150 mM NaCl, 2.5 mM KCl, 1 M MgCl_2_, and 1 M CaCl_2_). HEPES-buffered saline was poured over the gel until we initiated bioluminescence imaging. The level and areas of expression were confirmed by imaging Venus signals with a FVMPE-RS two-photon microscope (Olympus) and a Mai Tai DeepSee Ti:sapphire laser (Spectra-Physics) at 920 nm, through a 4x dry objective, 0.28 N.A. (Olympus) or a 25x water immersion objective, 1.05 N.A. (Olympus). Scanning and image acquisition were controlled using FV30S-SW image acquisition and processing software (Olympus).

For the bioluminescence imaging, a solvent of furimazine solution (Nano-Glo luciferase assay system, Promega) was evaporated with a VDR-20G vacuum desiccator (Jeio Tech) and a BSW-50N belt drive rotary vane vacuum pump (Sato Vac Inc.) overnight under dark conditions. The precipitate was eventually dissolved in propylene glycol (up to 5 mM). This solution was kept at −30 °C as a stock solution and used for further imaging experiments.

For *in vivo* imaging from awake animals, we attached a small o-ring (diameter: 10 mm) on the headplate using Kwik-Sil silicon adhesive (World Precision Instruments), as described in **Fig. 2d**. In some experiments, we alternatively used headplates consisting of an o-shaped window with a certain thickness (and two side bars to hold the mouse head)^34^. This circular pool area was used to keep the furimazine solution over the cranial window. During the bioluminescence imaging, the HEPES-buffered saline over the agarose gel was replaced with 50 μM furimazine solution (dissolved in 200 μl of the HEPES-buffered saline). This o-ring pool was eventually covered by a cover glass (15 mm diameter) glued over the o-ring with the Kwik-Sil adhesive. To minimize the reflection of the bioluminescence from LOTUS-V-expressing neurons at the outside of the brain, the surrounding areas, including surface of the headplate and dental cement, were stained black using Touch Up Paint X-1 matte black (SOFT99). This was important to suppress noisy signals from the target area. Further, the inside of the mouse chamber that was used for imaging during free movement was stained black using a black guard spray (Fine Chemical Japan).

### *In vivo* imaging of head-fixed mice

For signal detection, we used a Lumazone *in vivo* luminescence imaging system (Molecular Devices) equipped with Evolve Delta 512 EMCCD camera (Photometrics), C8600-05 GaAsP high-speed-gated image intensifier unit (Hamamatsu Photonics), W-VIEW GEMINI image splitting optics (Hamamatsu Photonics), and AT-X M100 PRO D macro lens (Tokina) for the recording. Although C8600-05 GaAsP image intensifier unit was used for other experiments, including the recordings from a primary culture of hippocampal neurons and freely-moving mice, C9546-02 GaAsP high-speed-gated image intensifier unit (Hamamatsu Photonics) equipped with a gate function was used to protect the image intensifier unit from the large current evoked by the visual stimulation light. Bioluminescence emitted from samples during imaging was split by a FF509-FDi01-25×36 dichroic mirror (Semrock) and passed through emission filters (NLuc channel; FF02-472/30-25 and FF01-483/32-25, Venus channel; FF01-537/26-25 and FF01-542/27-25) in W-VIEW GEMINI image splitting optics (Hamamatsu Photonics). During the recording from head-fixed mice, the mouse head was held using the attached headplate, and the mouse was placed over the custom-made running disk (**Fig. 2e**), which was attached to the rotary encoder to record the running speed of the mouse under the control of LabView (National Instruments). The cranial window was placed around the center of the field of view and the camera binning was set to 4.

For visual stimulation (light illumination to the right eye) during the bioluminescence imaging (from left V1 neurons in awake mice), a 15, 30, or 45-mW blue laser from Sapphire 458 LP (Coherent) was coupled to 3-mm-diameter liquid light guide (Thorlabs) through F35 plano-convex lens (Sigma), chopper wheel (Thorlabs), F40 plano-convex lens (Sigma), iris diaphragm (Sigma), and 12.7-mm-aperture optical shutter (Thorlabs), as shown in **Supplementary Fig. 3a**. A thin white tape covered the tip end of the liquid light guide to sufficiently scatter the output light and protect the mouse retina. The output power density of the 15, 30, and 45-mW blue laser was 0.55, 1.10, and 1.65 mW/cm^2^, respectively, at the surface level of the eye (there was a 1-cm distance between the eye and tip end of the liquid light guide).

During this experiment, an animal head was fixed under the camera lens by holding the attached headplate as described above. Three different light intensities were sequentially tested for each animal without changing the positions of the animal head and tip of the optical fiber. Light pulses (2-ms duration for a single pulse, with a 23-ms interval (40 Hz)) generated by a chopper wheel were delivered for 1 s, with an inter-trial interval of 7 s, using the optical shutter (**Supplementary Fig. 3b**). A total of 15 stimulus trials were repeated at each light intensity. Imaging was performed at 40 Hz, and the exposure time was set to 20 ms so that images were captured only when the light illumination was off (called “dead-time imaging”)^14,23^. Therefore, a light-driven artifact was not present in the detected signals. Exposure and gate-on timing were controlled using a TTL signal from a multifunction generator (WF1973, NF Corporation), with the output signal from the optical chopper system acting as the trigger.

### *In vivo* imaging of freely moving mice

During *in vivo* imaging, the freely moving animals were placed in the cage and signals were detected using the same equipment used for imaging head-fixed mice, except that wide-angle lens (HF12.5SA-1, Fujinon), C8600-05 GaAsP image intensifier unit (Hamamatsu Photonics) and no emission filters were used. The camera binning was set at 2 and 8 to record multiple mice and a single mouse, respectively. The frame rate was set at 10-100 Hz (they were adjusted according to the expression level and brightness of the bioluminescence to maintain a high signal-to-background ratio (SBR) and clearly identify the signal from the movie).

The merged movies shown in **Supplementary Video 1 and 3** were performed to confirm that the signal was correctly detected from the cranial window (**Supplementary Video 1**) or to capture the shapes of mice and target brain areas during imaging from multiple mice (**Supplementary Video 3**). These movies were produced using bright-field images acquired under a 632-nm Light Emitted Diode (LED) (LightEngine SPECTRA, Lumencor) illumination, and bioluminescence images acquired in the dark, which were captured alternately every 50 ms, as previously described^26,27^. The field of view was illuminated by LED with an irradiation time of 5 ms from the initiation of the camera exposure, once every two frames; the timing was controlled with a TTL signal from a multifunction generator (WF1973, NF Corporation). The TTL signal was generated using the output signal from the exposure time-out signals from an EMCCD camera as the trigger.

### Data analysis

All imaging data were processed using Fiji, R-software (Version 3.2.2.) and MATLAB (Mathworks). The curve fitting was performed using Origin 8.5.1 (OriginLab).

Neuronal activity was measured by calculating the FRET ratio (*R*) of LOTUS-V (Venus signal divided by NLuc signal) as previously described^14^. The background signal was subtracted from the signal of the specimen before the FRET ratio was calculated. The method of background subtraction depended on each imaging setup. For the cultured hippocampal neurons and head-fixed animal imaging, the “background signal” in both NLuc and Venus channels, measured in a randomly chosen region of interest (ROI) and placed at a non-specimen area, was subtracted from the signal of the specimen in each channel. When the freely moving animals were imaged, the “background image,” obtained by averaging 1000 blank images taken with the closed shutter of an EMCCD camera, was subtracted from the acquired images. The change in the FRET ratio (Δ*R*/*R*_0_) was calculated by subtracting the average value in the baseline (*R*_0_) from individual raw values at each time-point (Δ*R*=*Rt - R*_0_; *Rt* is a raw value at time-point “t”) and further dividing the difference, Δ*R*, by the *R*_0_. The meaning of the term “baseline” also depended on the experiment or analysis. During *in vitro* experiments in cultured hippocampal neurons, the term “baseline” meant that neurons were at the resting membrane voltage. The term “baseline” for locomotion analysis, however, meant that the animals were immobile and not moving (resting state, <5 cm/s for head-fixed mice, <1 cm/s for tracking data from freely-moving mice; the threshold of each was set based on the level of noise fluctuation of the detected locomotion velocity when mice were actually not moving, and by referring to a previous study^21^). The term “baseline” for the interaction analysis meant that an animal was distant enough from others (distant state, >4 cm) or in the “distant and resting” state, while for the visual stimulation trials, it meant that an animal was in the resting state during the absence of stimulation.

Each single imaging frame acquired via the W-VIEW GEMINI image splitting optics (Hamamatsu Photonics) contained information from two channels (Venus and NLuc signals), at either the left or right side. To align the two-channel data during post-processing, we first acquired non-bioluminescence images under white light so that the shape of the neuron (*in vitro*), cranial window (head-fixed mice), or cage (freely moving mice) could be similarly observed in each channel. These reference frames were separated to each channel. Then, using the coordinate information, the two channels of the actual imaging movies were aligned so that we could calculate the LOTUS-V FRET ratio (R) using common ROIs. During the *in vitro* and head-fix experiments, drifting or motion-based shift of the sample was rarely observed. However, to confirm that the ROI could correctly work in all frames of each movie, we performed motion correction as previously described^9^.

When we analyzed movies of primary culture of hippocampal neurons, the ROI was created from a mask image, which was made based on an averaged picture over all frames of each movie (with single-channel data). The threshold for bioluminescence intensity was manually determined to cover the expression area (e.g., top 0.5%; however, we obtained similar results using a variety of thresholds). The following Boltzmann function was used to fit the voltage-sensitive curve in primary hippocampal neuronal cultures:

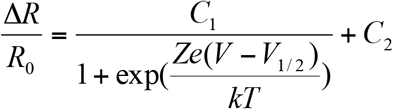

where, *C*_1_ and *C*_2_ were constant values, *e* was the elementary electric charge, *Z* was the effective valence, *k* was the Boltzmann constant, *T* was the room temperature in K, and *V*_1/2_ was the voltage at which Δ*R*/*R*_0_ is half-activated.

To analyze voltage kinetics, we fitted the activation and deactivation curves of Δ*R*/*R*_0_ using the following two-component exponential equation:

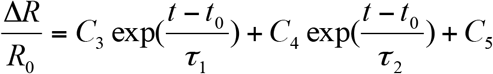

where, *C*_3_, *C*_4_, and *C*_5_ were constant values (*C*_3_ and *C*_4_ were used to calculate the fraction of *τ*_fast_), *t*_0_ was the initial time-point, and *τ*_1_ and *τ*_2_ were the time constants for the fast and slow components.

The protocol used for *in vitro* experiments were also used to create the ROIs when analyzing the data from the movies of head-fixed mice. The relationship between the LOTUS-V signal and mouse locomotion was analyzed as described above.

When imaging the freely moving mice, the bioluminescence spot derived from the cranial window in the field of view was automatically tracked using the “Particle Track Analysis” (PTA ver 1.2; https://github.com/arayoshipta/projectPTAj). Further, the cranial window in the headplate seen in the bright field image was manually tracked using “Manual Tracking” (ImageJ plugin) when analyzing interactively locomoting mice. The trajectories shown in **Fig. 3b, Fig. 4a, and Supplementary Videos 2 and 4,** were pseudo-colored, based on the length of the line segments or z-normalized Δ*R*/*R*_0_.

For automatic tracking, movies during bioluminescence imaging from freely moving mice were processed using a Gaussian blur filter with a radius value set to 1.0. Each movie was separated into the NLuc and Venus channels, referring to the bright field image as mentioned above. The separated movies were processed using the “AND” function of Image J to produce the reference movie containing information about the intensity of both channels. This reference movie was only used to obtain the coordinate information of a bioluminescence spot by PTA, which was advantageous to increase the SBR and more accurately detect its position. Thereafter, the bioluminescence intensity in each channel was measured based on this information. Since the signal intensity (not the FRET ratio) was affected by the angle of the mouse head, some of the frames were difficult to analyze automatically due to the low SBR. Therefore, when the signal could not be automatically detected over 3 frames using PTA, these missed frames were excluded from the analysis. If the signal appeared within 3 frames (i.e., with two frames missed at most) from the position where the last signal was detected, then the PTA recognized them as a series, placed the ROI at the same position, and continued automatic tracking. In these cases, linear interpolation was used to fill the missing values.

Due to the fluctuation of background noise, the signal intensity after the background subtraction was sometimes lower than 0, which is also often observed in single molecule analysis^35^. This negative value was almost 0 but could disperse the distribution of ratio value. Thus, the absolute value of the minimum value was added to the all values in both the NLuc and Venus channels. If the signal intensity in the NLuc channel was 0, the frame at that time-point was removed before data analysis. Shot noise during imaging was main and strong source of artifacts (0~0.8), which was sometimes much stronger than the actual signals (>2.0), and often overlapped with the bioluminescence signal in the image. To minimize the effect of the shot noise, ratios within the 0.8-2.0 range was selectively used for data analysis. This range was carefully set by referring to the result of the head-fix mouse (the averaged ratio value plus or minus threefold the value of standard deviation, N=8 animals). Locomotion velocity was calculated from the frame-by-frame position changes in the center of gravity of the bioluminescence signal derived from the mouse brains.

The SBR was calculated by dividing the averaged intensity of NLuc and Venus emissions by the background intensity. Automatic tracking could not properly distinguish the bioluminescence signal from shot noise when the SBR was lower than 0.12 (due to substrate consumption and/or an original low-expression level of LOTUS-V). Thus, low SBR data were excluded from the analysis. The total imaging period for each animal was estimated by the duration of high SBR (0.12 or higher).

When imaging multiple mice, 1 mouse (of the 4 mice) was relatively immobile and did not voluntarily interact with others (**Fig. 4, Supplementary Video. 3-4**). To quantitatively evaluate how immobile each mouse was, the fraction of the active state (>1 cm/s)^21^ was calculated. Although the average fraction of the mice used for the free-moving experiments was 64.2±7.8% [mean ± SE.] (N=11 animals), that of the quiet one (N=1) was only 2.1%. Following justification of the normal distribution of the data (Kolmogorov-Smirnov test, p >0.05, N=12 animals), a “Chi-squared test for outliers” systematically detected this quiet animal as an outlier (p <0.05). Therefore, the data of this mouse was excluded from the analysis.

To statistically compare results obtained from different animals expressing either LOTUS-V or LOTUS-V(D129R), the Δ*R*/*R*_0_ in each animal was z-normalized^36^ to obtain “z-normalized *ΔR/R_0_”*. The z-score values were calculated by subtracting the average baseline signals from individual raw values, and further dividing the difference by the baseline standard deviation. When we analyzed the data during visual stimulation, first 0.5 s of data was used as the signals during visual stimulation, while baseline (resting state) signals were calculated by randomly choosing the same number of frames (with that of data during visual stimulation) from whole resting state data of each animal for respective comparisons (i.e., 10,000 repeats of random choices were used to calculate the representative (average) value for each “animal,” at each light intensity; or one time random selection to obtain a group of baseline signals for each “trial” at each light intensity).

To analyze the activity of V1 during the interaction of multiple mice, the area of each mouse in the 2D image was approximated using three circles (**Supplementary Fig. 4a**), by referring to a previous study^37^. The positions of the nose (x_1_, y_1_), headplate (x_2_, y_2_), and tail root (x_3_, y_3_) were manually tracked. This coordinate information was further used to calculate the positions of the cervix (2x_2−_x_1_, 2y_2−_y_1_) and dorsum ((x_3_ + 2x_2_ − x_1_)/2, (y_3_ + 2y_2_ − y_1_)/2). The centers of three variable circles were then set at the headplate (Circle 1), cervix (Circle 2), and dorsum (Circle 3), respectively. The radius of each circle was set as the distance between the headplate and nose (for Circle 1 and Circle 2), or that between the dorsum and tail root (for Circle 3). Thereafter, the distances between the nose of the target animal and edge of each circle were calculated. The shortest one was used as the index of the interaction analysis (**Supplementary Fig. 4b and c**). When the nose was the inside of the circle, the distance was represented as a negative value.

Since LOTUS-V reported an increase of V1 activity by visual stimulation (**Supplementary Fig. 3c**), one might be concerned that bioluminescence from other animals could work as visual input. To exclude this possibility, we evaluated the power density of LOTUS-V bioluminescence from the V1 surface. The number of photons, *P,* collected on an EMCCD camera sensor was calculated from the total analogue-to-digital converter (ADC) counts, I, using the following equation:

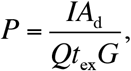

where, *A*^d^ was the analog to digital conversion factor of an EMCCD camera, *Q* was the quantum efficiency of an EMCCD camera, *t*_ex_ was the exposure time, and *G* was the radiant emittance gain of an image intensifier unit. Since the ray divergence, *θ*, from the bioluminescent object was small enough, the fraction of the collected photons, *F*, was calculated using the following equation:

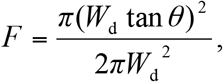

where *W*_d_ was the working distance of a lens.

Therefore, the bioluminescence intensity, *B*, radially emitted from the V1 surface was calculated using the following equation:

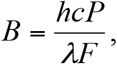

where, *h* was the Planck constant, *c* was the speed of light; and *λ* was the wavelength. Accordingly, the power density of LOTUS-V bioluminescence from V1 surface was computed to be (7.6±1.0) × 10^−9^ and (9.0±1.3) × 10^−9^ mW/cm^2^ (N=6 animals), at 480 nm and 540 nm, respectively. Since the weakest visual stimulation (0.55 mW/cm^2^) did not influence the LOTUS-V signal (**Supplementary Fig. 3c**), the increase of V1 activity during the interaction was unlikely to be caused by the bioluminescence from other animals (**Fig. 4c-d**).

### Statistical Analyses

Statistical analyses were conducted using R software or MATLAB. The type of the statistical analysis conducted for each analysis was described in the manuscript. All p-values less than 0.0001 were described as “P<0.0001.” All statistical tests, except the Chi-squared test, were performed as “two-tailed” tests. Statistical significance was set at P <0.05.

## References

1. Marshall, J. D. et al. Cell-Type-Specific Optical Recording of Membrane Voltage Dynamics in Freely Moving Mice. Cell 167, 1650–1662.e15 (2016).

2. Buzsáki, G., Anastassiou, C. & Koch, C. The origin of extracellular fields and currents--EEG, ECoG, LFP and spikes. Nat. Rev. Neurosci. 13, 407–420 (2012).

3. Mohajerani, M. H. et al. Spontaneous cortical activity alternates between motifs defined by regional axonal projections. Nat. Neurosci. 16, 1426–1435 (2013).

4. Akemann, W., Mutoh, H., Perron, A., Rossier, J. & Knöpfel, T. Imaging brain electric signals with genetically targeted voltage-sensitive fluorescent proteins. Nat. Methods 7, 643–649 (2010).

5. Akemann, W. et al. Imaging neural circuit dynamics with a voltage-sensitive fluorescent protein. J. Neurophysiol. 108, 2323–2337 (2012).

6. Gong, Y. et al. High-speed recording of neural spikes in awake mice and flies with a fluorescent voltage sensor. Science 350, 1361–1366 (2015).

7. Carandini, M. et al. Imaging the Awake Visual Cortex with a Genetically Encoded Voltage Indicator. J. Neurosci. 35, 53–63 (2015).

8. Chen, T. W. et al. Ultrasensitive fluorescent proteins for imaging neuronal activity. Nature 499, 295–300 (2013).

9. Agetsuma, M., Hamm, J. P., Tao, K., Fujisawa, S. & Yuste, R. parvalbumin-Positive Interneurons Regulate Neuronal Ensembles in Visual Cortex. Cereb. Cortex In press (2017).

10. Higashi A, Uchizono K, Tani Y, Hoshino M, Yano T, Y. K. Real time online data processing system for the e.e.g, and body movement during the lifetime of a freely moving mouse. Med. Biol. Eng. 17, 416–418 (1979).

11. Ferezou, I., Bolea, S. & Petersen, C. C. H. Visualizing the Cortical Representation of Whisker Touch: Voltage-Sensitive Dye Imaging in Freely Moving Mice. Neuron 50, 617–629 (2006).

12. Ziv, Y. & Ghosh, K. K. Miniature microscopes for large-scale imaging of neuronal activity in freely behaving rodents. Curr. Opin. Neurobiol. 32, 141–147 (2015).

13. Miyamoto, D. & Murayama, M. The fiber-optic imaging and manipulation of neural activity during animal behavior. Neurosci. Res. 103, 1–9 (2016).

14. Inagaki, S. et al. Genetically encoded bioluminescent voltage indicator for multi-purpose use in wide range of bioimaging. Sci. Rep. 7:42398, (2017).

15. Murata, Y., Iwasaki, H., Sasaki, M., Inaba, K. & Okamura, Y. Phosphoinositide phosphatase activity coupled to an intrinsic voltage sensor. Nature 435, 1239–1243 (2005).

16. Hall, M. P. et al. Engineered Luciferase Reporter from a Deep Sea Shrimp Utilizing a Novel Imidazopyrazinone Substrate. ACS Chem Biol. 7, 848–857 (2012).

17. Nagai, T. et al. A variant of yellow fluorescent protein with fast and efficient maturation for cell-biological applications. Nat. Biotechnol. 20, 87–90 (2002).

18. Chamberland, S. et al. Fast two-photon imaging of subcellular voltage dynamics in neuronal tissue with genetically encoded indicators. Elife 6, e25690 (2017).

19. Jin, L. et al. Single Action Potentials and Subthreshold Electrical Events Imaged in Neurons with a Fluorescent Protein Voltage Probe. Neuron 75, 779–785 (2012).

20. Keller, G. B., Bonhoeffer, T. & Hübener, M. Sensorimotor Mismatch Signals in Primary Visual Cortex of the Behaving Mouse. Neuron 74, 809–815 (2012).

21. Saleem, A. B., Ayaz, A., Jeffery, K. J., Harris, K. D. & Carandini, M. Integration of visual motion and locomotion in mouse visual cortex. Nat. Neurosci. 16, 1864–1869 (2013).

22. Fu, Y. et al. A Cortical Circuit for Gain Control by Behavioral State. Cell 156, 1139–1152 (2014).

23. Chang, Y. F., Arai, Y. & Nagai, T. Optogenetic activation during detector ‘ dead time’ enables compatible real-time fluorescence imaging. Neurosci. Res. 73, 341–347 (2012).

24. Niell, C. M. & Stryker, M. P. Modulation of Visual Responses by Behavioral State in Mouse Visual Cortex. Neuron 65, 472–479 (2010).

25. Hamada, T. et al. In vivo imaging of clock gene expression in multiple tissues of freely moving mice. Nat. Commun. 7, 11705 (2016).

26. Saito, K. et al. Luminescent proteins for high-speed single-cell and whole-body imaging. Nat. Commun. 3, 1262 (2012).

27. Matsushita, J. et al. Fluorescence and Bioluminescence Imaging of Angiogenesis in Flk1-Nano-lantern Transgenic Mice. Sci. Rep. 7, 46597 (2017).

28. Madisen, L. et al. Transgenic Mice for Intersectional Targeting of Neural Sensors and Effectors with High Specificity and Performance. Neuron 85, 942–958 (2015).

29. Wheeler, M. A. et al. Genetically targeted magnetic control of the nervous system. Nat Neurosci 19, 756–761 (2016).

30. Montgomery, K. L. et al. Wirelessly powered, fully internal optogenetics for brain, spinal and peripheral circuits in mice. Nat. Methods 12, 969–974 (2015).

## References

31. Zhao, Y. et al. Microfluidic cell sorter-aided directed evolution of a protein-based calcium ion indicator with an inverted fluorescent response. Integr. Biol. (Camb). 6, 714–725 (2014).

32. Inutsuka, A. et al. Concurrent and robust regulation of feeding behaviors and metabolism by orexin neurons. Neuropharmacology 85, 451–460 (2014).

33. Grimm, D. et al. In Vitro and In Vivo Gene Therapy Vector Evolution via Multispecies Interbreeding and Retargeting of Adeno-Associated Viruses. J. Virol. 82, 5887–5911 (2008).

34. Goldey, G. J. et al. Removable cranial windows for long-term imaging in awake mice. Nat. Protoc. 9, 2515–2538 (2014).

35. Wang, Y. et al. Single molecule FRET reveals pore size and opening mechanism of a mechano-sensitive ion channel. Elife 3, e01834 (2014).

36. Herry, C. et al. Switching on and off fear by distinct neuronal circuits. Nature 454, 600–606 (2008).

37. de Chaumont, F. et al. Computerized video analysis of social interactions in mice. Nat. Methods 9, 410–417 (2012).

